# The transferable resistome of produce

**DOI:** 10.1101/350629

**Authors:** Khald Blau, Antje Bettermann, Sven Jechalke, Eva Fornefeld, Yann Vanrobaeys, Thibault Stalder, Eva Top, Kornelia Smalla

## Abstract

Produce is increasingly recognized as a reservoir of human pathogens and transferable antibiotic resistance genes. This study aimed to explore methods to characterize the transferable resistome of bacteria associated with produce. Mixed salad, arugula, and cilantro purchased from supermarkets were analyzed by means of cultivation- and DNA-based methods. Before and after a nonselective enrichment step, tetracycline (tet) resistant *Escherichia coli* were isolated and plasmids conferring tet resistance were captured by exogenous plasmid isolation. Tet resistant *E. coli* isolates, transconjugants and total community (TC)-DNA from the microbial fraction detached from leaves or after enrichment were analyzed for the presence of resistance genes, class 1 integrons and various plasmids by real-time PCR and PCR-Southern blot hybridization. Real-time PCR primers were developed for IncI and IncF plasmids. Tet resistant *E. coli* isolated from arugula and cilantro carried IncF, IncI1, IncN, IncH11, IncU and IncX1 plasmids. Three isolates from cilantro were positive for IncN plasmids and *bla*_CTX-M-1_. From mixed salad and cilantro, IncF, Inc11, and IncP-1β plasmids were captured exogenously. Importantly, whereas direct detection of IncI and IncF plasmids in TC-DNA failed, these plasmids became detectable in DNA extracted from enrichment cultures. This confirms that cultivation-independent DNA-based methods are not always sufficiently sensitive to detect the transferable resistome in the rare microbiome. In summary, this study showed that an impressive diversity of self-transmissible multiple resistance plasmids was detected in bacteria associated with produce that is consumed raw, and exogenous capturing into *E. coli* suggests that they could transfer to gut bacteria as well.

## IMPORTANCE

Produce is one of the most popular food commodities. Unfortunately, leafy greens can be a reservoir of transferable antibiotic resistance genes. We found that IncF and IncI plasmids were the most prevalent plasmid types in *E. coli* isolates from produce. This study highlights the importance of the rare microbiome associated with produce as a source of antibiotic resistance genes that might escape cultivation-independent detection, yet may be transferred to human pathogens or commensals.

## INTRODUCTION

Despite its benefit to human health, consumption of produce is increasingly recognized as a source of pathogenic bacteria, antibiotic-resistant bacteria (ARB) and antibiotic resistance genes (ARGs) associated with mobile genetic elements (MGEs) (1–5). Recently, several food-borne disease outbreaks have been associated with produce contamination world-wide (5–9). The microbiome of produce is important for plant health and vigor and was shown to be highly dynamic during growth, post-harvest, and processing (10), but can also contain potentially pathogenic bacteria from human and animals sources, including *E. coli* strains (11). Contamination can occur pre-harvest (i.e., through organic fertilizers, soil, wild animals or contaminated irrigation water) and post-harvest during processing (12, 13).

Antibiotic resistance in bacterial pathogens has increased globally due to the widespread use and misuse of antibiotics (14–17). Antibiotic resistance levels in *E. coli* are useful indicators of overall resistance levels of bacteria on foods, in animals, and humans (11). Antibiotic resistance and ARGs have been documented for enteric bacteria from various produce, which could facilitate the dissemination of resistant bacteria to a wider community of people (1, 2, 4, 16, 18, 19). If ARGs are localized on MGEs such as plasmids or conjugative transposons they can be transferred horizontally to pathogens (20). Horizontal gene transfer (HGT) takes place at sites with high cell densities of plasmid donors and recipients, nutrient availability and selective pressure. The phytosphere, including the rhizosphere and the phyllosphere have been reported as hot spots of HGT (21). The plasmid-mediated resistome of produce bacteria might provide the enterobacteria with ARGs in the intestine under selective conditions.

Conjugative plasmids can often confer not only resistance towards multiple antibiotics but also towards heavy metal compounds or disinfectants making co-selection possible (22–25). Although plasmids belonging to the incompatibility groups IncF and IncI have a narrow host range (NHR), they are assumed to be important for the dissemination of ARGs in *E. coli* and other *Enterobacteriaceae* (26, 27). Most importantly, resistance and virulence-associated traits of *E. coli* isolates were almost exclusively found on IncF group plasmids (28, 29). However, no real time (RT-) PCR systems that allow the cultivation-independent detection and quantification of these plasmids in total community (TC)-DNA are available.

In this study, we employed a combination of culture-dependent and -independent approaches to assess the transferable resistome of bacteria associated with produce (see Figure 1). Tetracycline (tet) resistant *E. coli* were isolated from produce directly after purchase and after seven days of storage by selective plating with and without prior non-selective enrichment. In addition, we captured a set of transferable tet resistance plasmids into a *E. coli* recipient strains using the so-called exogenous plasmid isolation method (30). Tetracycline was chosen because of its wide use in animal husbandries resulting in a high load released via organic fertilizers into the agro-ecosystem (31). TC-DNA was also extracted from the microbial fraction detached from produce or after nonselective enrichment to detect and quantify the abundance of ARGs and MGEs. New RT-PCR primers were developed for the detection and quantification of IncF and IncI plasmids. Our study showed the presence of diverse transferable antibiotic resistance plasmids in the rare microbiome of produce, which would have likely been missed by exclusively using metagenomic approaches.

**Figure 1.**
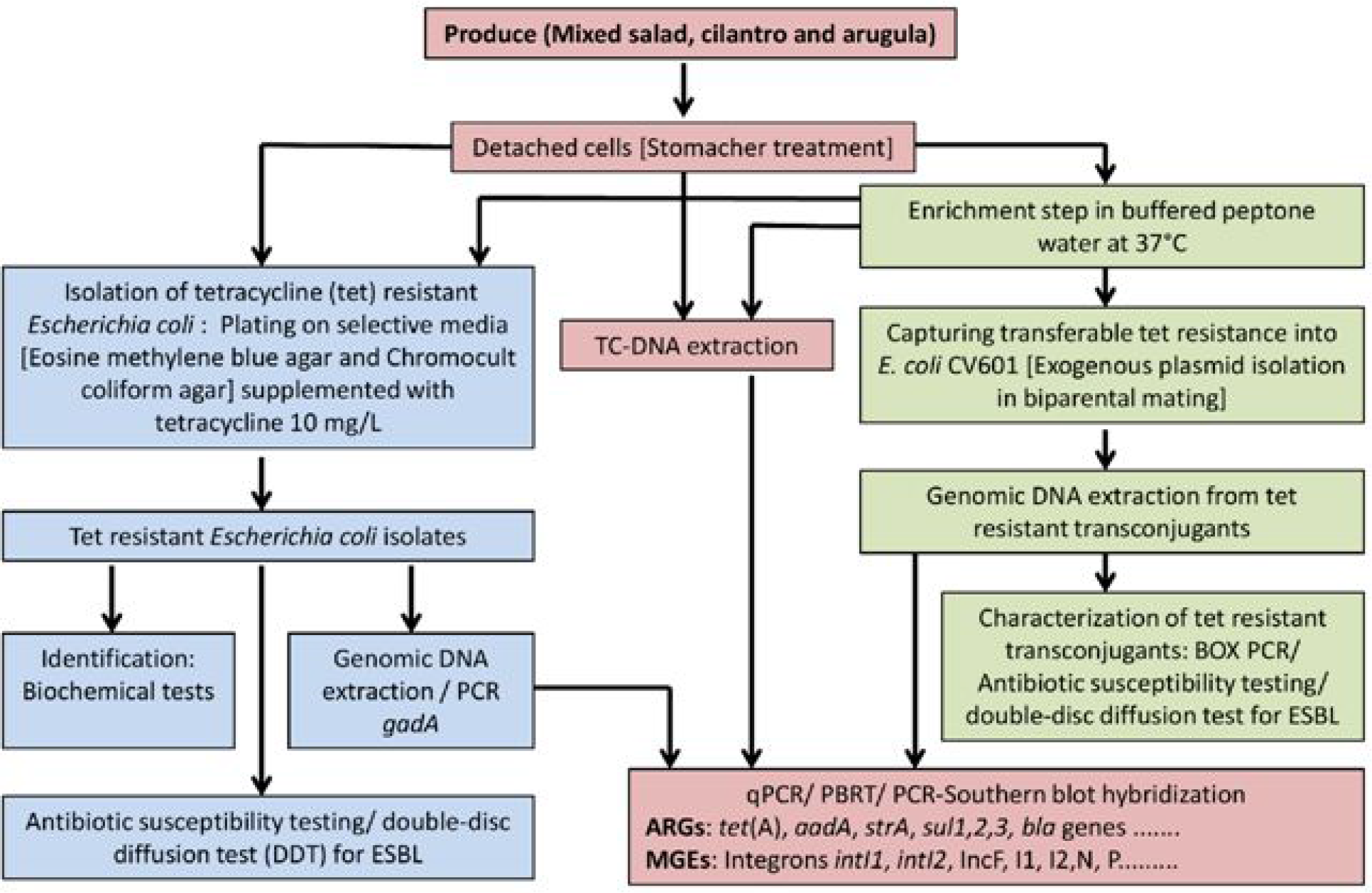
Flow diagram of the experimental set-up of this study to employ culture-dependent and -independent approaches to characterize the transferable resistome of bacteria associated with produce.

## RESULTS

### Phenotypic and genotypic characterization of tet resistant *E. coli* isolates

To determine if produce was a source of antibiotic resistant *E. coli*, we determined the occurrence and resistance profiles of tet resistant *E. coli* isolated from 24 samples of produce directly or after an overnight enrichment step. The characterization of a total of 63 tet resistant *E. coli* isolates from cilantro (n=54), arugula (n=7) and mixed salad (n=2), of which 50 were recovered after nonselective enrichment and 13 without enrichment (20.6%) revealed an impressive diversity (Table 1).

**TABLE 1.**
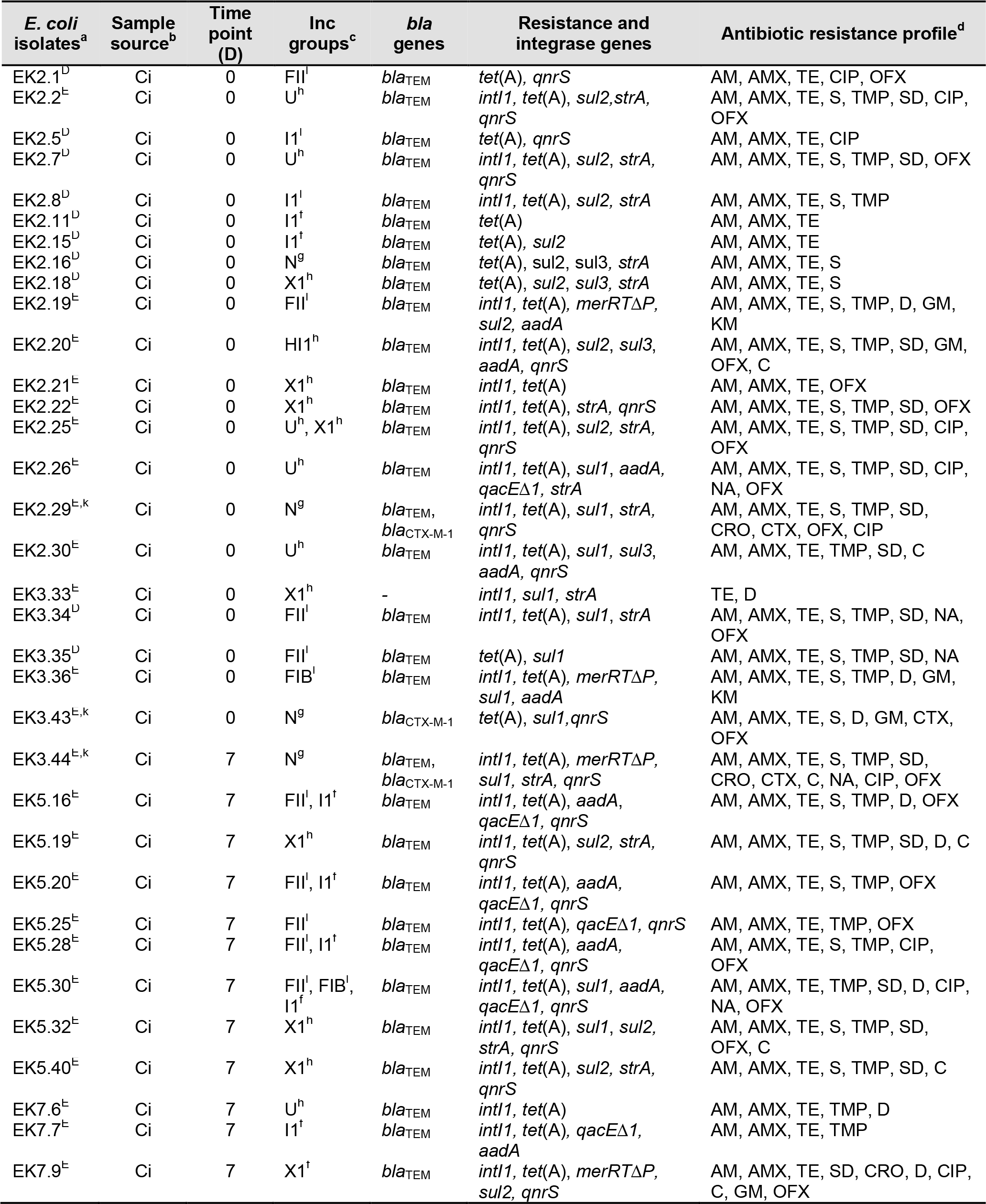
Characterization of representative tetracycline resistant *E. coli* isolates from fresh produce

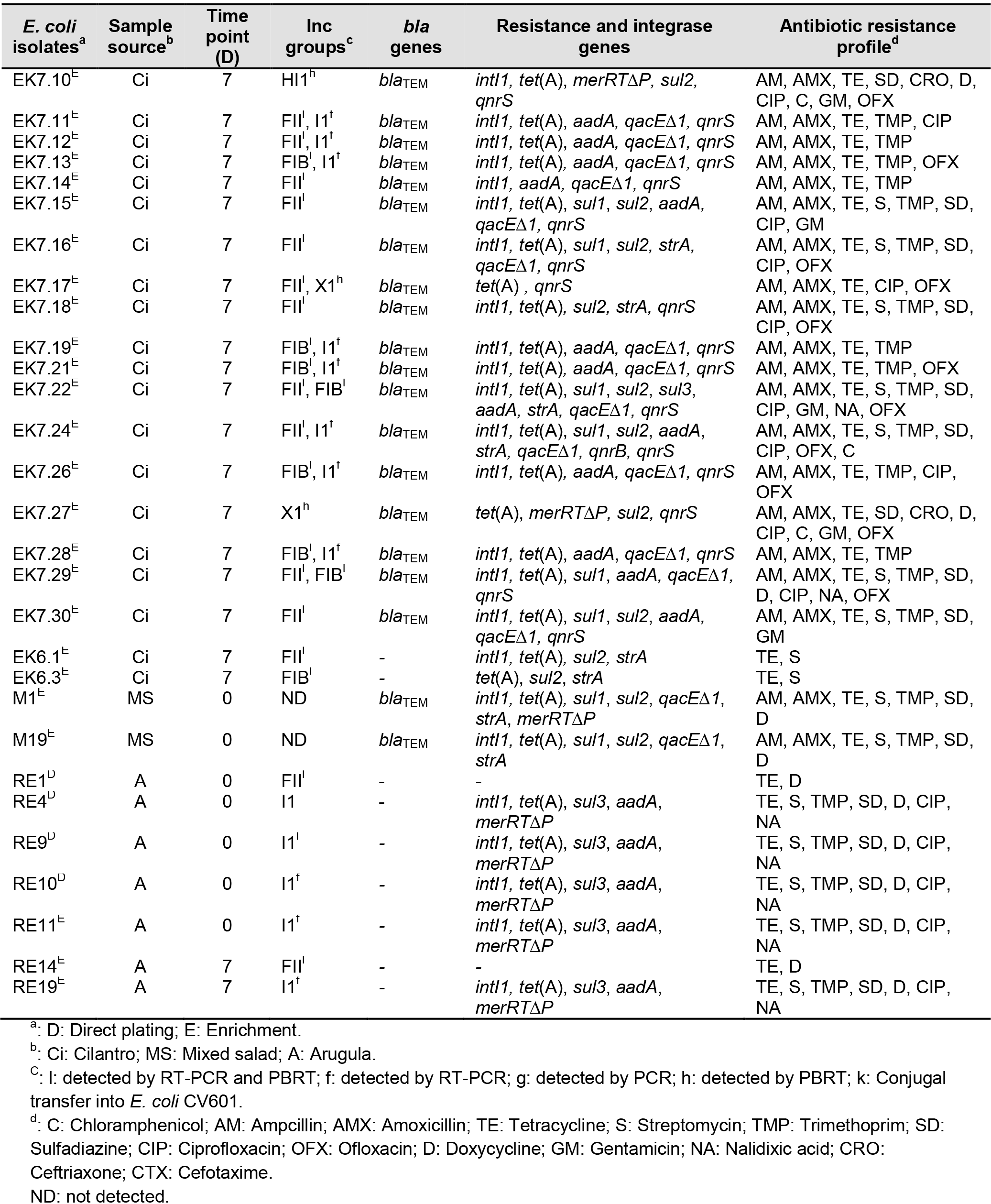

Almost all *E. coli* isolates were resistant to antibiotics from at least one class, and two isolates were resistant to eight antibiotic classes: Tetracyclines (TE; D), penicillins (AM;AMX), 3^rd^ generation cephalosporins (CTX; CRO), fluoroquinolones (CIP; OFX; NA), aminoglycosides (GM; S), sulfonamides (SD), phenicols (C), and trimethoprim (TMP). Most of the tet resistant *E. coli* displayed in addition resistance to ampicillin and amoxicillin (84%), followed by trimethoprim (73%). Resistances to ofloxacin, ciprofloxacin, sulfadiazine, and streptomycin were also common. We tested all the isolates for the production of extended-spectrum beta-lactamases (ESBLs) with the double-disc diffusion assay (DDT) and found three ESBL-producing *E. coli* which were isolated from two of the cilantro samples.

We then tested the genomic DNA from the collection of *E. coli* isolates for the presence of various resistance genes [*tet*(A), *strA, sul1, sul2, sul3, aadA, qacE/qacEΔ1, merRTΔP, bla* genes (TEM, CTX-M, and SHV), *qnr* genes (*qnr*A, *qnr*B, and *qnr*S)], and integrase genes *intI1* and *intI2* by RT- or regular PCR (Table 1). The most commonly detected ARGs was the tetracycline resistance gene *tet*(A) in 59 out of 63 isolates. A total of 10 isolates were positive for sulfonamide resistance genes *sul1*, 14 for *sul2* and five for *sul3*. The combination of *sul1-sul2, sul2-sul3*, and *sul1-sul3* were detected in seven, three and one isolates, respectively. All three *sul* genes were found in one tet resistant *E. coli* isolate from cilantro. *qnrB* and *qnrS* genes encoding fluoroquinolone resistance were detected alone or in combination in one and 38 isolates, respectively. The *bla*_TEM_ genes encoding for the resistance to ampicillin and amoxicillin were detected in 82.5% of tet resistant *E. coli* isolates. The *bla*_CTX-M-1_ gene encoding ESBL resistance was only detected in three isolates and was found in combination with *bla*_TEM_ genes in two *E. coli* from cilantro. The *bla*_SHV_ gene encoding ESBL resistance was not detected in any of the isolates. For streptomycin/spectinomycin resistance gene, *aadA* (24 isolates) was most common, followed by *strA* (21 isolates) and *aadA-strA* (three isolates).

The class 1 integron integrase gene *intI1* was detected in 50 isolates, while the class 2 integron integrase gene *intI2* was not detected at all. Although *qacE/qacEΔ1* encoding quaternary ammonium compound resistance is a typical component of class 1 integrons, the gene was detected only in 23 isolates, suggesting a large proportion of atypical class 1 integrons. Interestingly, *merRTΔP* encoding for regulatory, transport, and extracellular mercury-binding genes was detected in 12 isolates. These findings show that produce can be a source of multidrug resistant *E. coli* isolates.

### Characterization of plasmids in tet resistant *E. coli* isolates

To test if the tet resistant *E. coli* isolates recovered from produce harbor plasmids and to assign them to known plasmid groups, their genomic DNA was screened by TaqMan probe-based RT-PCR systems for IncF and IncI plasmids and by PCR-based replicon typing (PBRT) (Table 1). All isolates that were positive by RT-PCR targeting the IncF (*traI* gene) were also identified by replicon typing as IncF confirming the specificity of the novel TaqMan RT-PCR system. However, PBRT allowed also the assignment to the different IncF subgroups. Furthermore, other plasmids were also identified by PBRT or RT-PCR (*korB*, specific for IncP-1 plasmids) or PCR (IncN). A summary of the plasmid/replicon types detected among the 63 representative tet resistant *E. coli* isolates is given in Table 1. For cilantro and arugula, almost all tet resistant *E. coli* isolates contained plasmids [61 out of 63], but the plasmids detected in the two isolates from mixed salad could not be assigned using the RT-PCR or PBRT. In most isolates (n=45) one plasmid type was detected, but some had two (n=15) or three (n=1) plasmids. Plasmids from seven different Inc groups were found in the 63 *E. coli* isolates: IncFII (n=21), IncI1 (n=17), IncX1 (n=11), IncFIB (n=10), IncU (n=6), IncN (n=4), and IncHI1 (n=2). All Inc groups were found in *E. coli* isolates from cilantro, whereas arugula resulted in only two groups [IncI1 (n=5) and IncFII (n=2)]. Plasmids of the IncF groups (FII and FIB) were the predominant types, followed by IncI1 and IncX1 plasmids. The combination of replicon types IncFII and IncFIB was detected in two isolates, whereas the combination of replicon types IncFII-IncI1 and IncFIB-IncI1 was found in six and five isolates, respectively. In one isolate from cilantro the combination IncFII-IncFIB-IncI1 was detected. IncI2 plasmids were not detected in any of *E. coli* isolates.

### Conjugal transfer of antibiotic resistance

Conjugation experiments were conducted in order to determine the potential transfer of antibiotic resistances to other bacteria. Conjugal transfer experiments were performed using tet resistant *E. coli* isolates positive for ESBL (EK2.29, EK3.43, and EK3.44) as donors and kanamycin and rifampicin resistant *E. coli* CV601 as a recipient at 37°C. We selected transconjugants on LB plates containing tetracycline and cefotaxime, which corresponded to phenotypes of the donors. The transfer of the resistance phenotypes was successful.

### Phenotypic and genotypic characterization of plasmids captured via exogenous isolation

We further investigated the presence of transferable plasmids in produce by capturing tet resistance plasmids from nonselective enrichment culture of fresh leaves from cilantro, mixed salad, and arugula by exogenous plasmid isolation into *E. coli* CV601. Tet resistant transconjugants were only captured on day 0 but not on day 7. The transfer frequencies of tet resistant transconjugants were 1.73 × 10 ^−7^, 1.55 × 10^−4^ and 4.66 × 10^−9^ per recipient in cilantro, mixed salad, and arugula, respectively. While all transconjugants obtained from cilantro (n=27) and arugula (n=23) were characterized, from mixed salad only a total of 41 transconjugants was analyzed due to the high number of transconjugants obtained. Based on an initial phenotypic and genotypic analyses, 15 representative out of 91 tet resistant transconjugants from produce (cilantro, n=12; arugula, n=1; mixed salad, n=2) were selected for further characterization. The majority of these transconjugants acquired resistance to at least two antibiotic classes, and all were resistant to tetracycline, ampicillin and amoxicillin.

The *bla*_TEM_ genes encoding for ampicillin and amoxicillin resistances were detected in 86.7% of tet resistant transconjugants (Table 2). The tetracycline resistance gene *tet*(A) was found in 13 out of 15 transconjugants from cilantro and arugula but not from mixed salad, while *tet*(Q) was only detected in one plasmid (pBMS1) isolated from the mixed salad. Four tetracycline resistance plasmids (pBC1.1, pBC1.3, pBC1.9, and pBC1.12) captured from cilantro carried the insertion sequence IS *1071* and class 1 integrons (*intI1*), tetracycline resistance gene *tet*(A), but encoded also resistance to ampicillin (*bla*_TEM_), and mercury compounds (*merRTΔP*). Eight plasmids from cilantro transconjugants (pBC2.1, pBC2.2, pBC2.3, pBC2.4, pBC2.6, pBC2.8, pBC2.11, and pBC2.15) carried *tet*(A), *qnrS*, and *bla*_TEM_ and two of them (pBC2.1 and pBC2.4) carried in addition *sul1* and sul2, respectively. Two tet resistance plasmids (pBMS1 and pBMS4) captured from mixed salad carried *sul1, strA, merRTΔP*, *bla*_TEM_, and *intI1*. One plasmid (pBA1) captured from arugula carried *bla*_TEM_ and *tet*(A) (Table 2). Thus this approach demonstrates that transferable multidrug resistance plasmids were easily captured by *E. coli* CV601, a process that might also occur in the human gut.

**TABLE 2.**
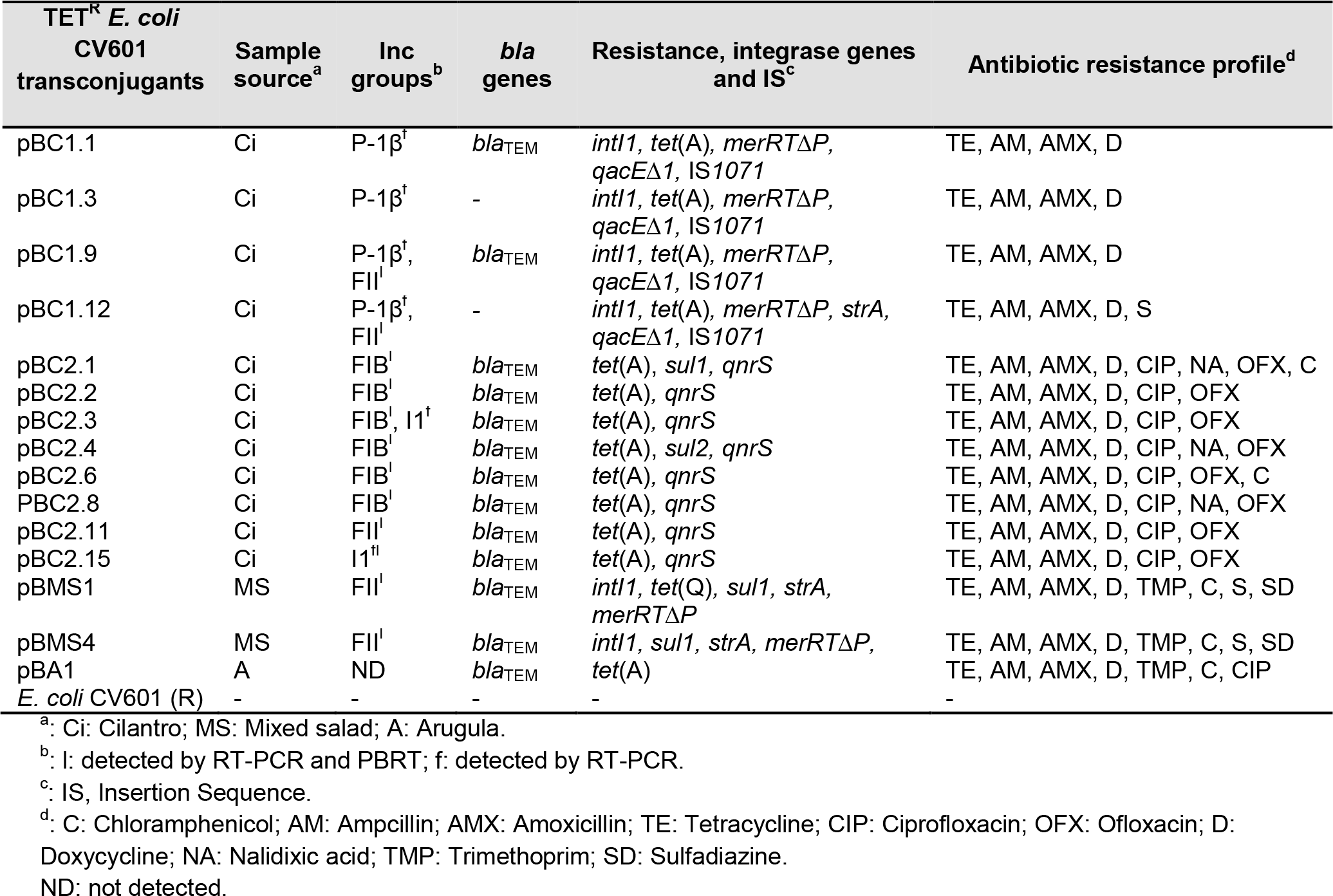
Characterization of representative tetracycline resistant *E. coli* CV601 transconjugants captured from produce.

### Identification of exogenously isolated plasmids

The newly developed TaqMan probe-based RT-PCR assay was used to screen the tet resistant transconjugants for the presence of IncF and IncI plasmids and validated by PBRT. In addition, other plasmids were also identified by RT-PCR (*korB*, specific for IncP-1 plasmids) and Southern blot hybridization. Plasmids of known Inc groups were detected in all transconjugants from the mixed salad and cilantro but not in the transconjugants from arugula. Representative transconjugants from cilantro and mixed salad carried either one (n=11) or two (n=3) replicons. In 12 transconjugants from cilantro samples four different plasmid replicon types were detected (Table 2): IncFII (n=3); IncFIB (n=6); IncI1 (n=2); and IncP-1β (n=4). In contrast, the transconjugants isolated from mixed salad showed only one replicon type, IncFII (n=2). One plasmid that could not be assigned by PBRT or RT-PCR was isolated from arugula leaves. The combination of replicon types IncFII and IncP-1β was detected in two transconjugants (pBC1.9 and pBC1.12), while the combination of replicon types of plasmids IncFIB and Inc11 was found in one transconjugant (pBC2.3) captured from cilantro leaves. Southern blot hybridization for sequences specific for IncP-1 plasmids revealed that four plasmids belonged to the IncP-1β subgroup. IncI2 plasmids were not detected in any tet resistant transconjugants (Table 2). In contrast to IncFIB/FII and IncI1 plasmids, the IncP-1β plasmids captured exogenously were not detected in the 63 tet resistant *E. coli* isolates.

### Detection of IncF and IncI plasmids, *tet*(A), and *intl1* in total community DNA

We also screened for plasmids (IncF, Inc11 and, IncI2), tetracycline resistance genes *tet*(A) and integrase gene *intI1* in TC-DNA extracted from bacterial communities either directly after their detachment from fresh leaves or after an enrichment step, using PCR-Southern blot hybridization and RT-PCR (Table 3). Using the RT-PCR method, IncF and IncI plasmids as well as *tet*(A) gene were detected in TC-DNA extracted from enrichment cultures of leaves, but not in TC-DNA from the detached bacteria. In contrast, the *intI1* gene was detected in both kinds of TC-DNA. Consistent with these results, PCR-Southern blot hybridization targeting the IncF and IncI plasmids and *tet*(A) revealed strong hybridization signals in TC-DNA extracted from the enrichment cultures, but very weak or no signals from direct extractions.

**TABLE 3.** PCR hybridization and real-time PCR of IncF, I1, I2 plasmids and *intI1* and *tet*(A) from TC-DNA extracted from fresh produce before and after enrichment.

| Fresh produce | DNA Isolation | Time point (D) | IncF |  | IncI1 |  | IncI2 |  | *int1* |  | *tet(A)* |  |
| --- | --- | --- | --- | --- | --- | --- | --- | --- | --- | --- | --- | --- |
|  |  |  | qPCR | Blot | qPCR | Blot | qPCR | Blot | qPCR | Blot | qPCR | Blot |
| Mixed salad | Direct extraction | 0 | - | - | - | - | - | +++ | + | (+++) | - | (++)<sup>1*</sup> |
|  |  | 7 | - | - | - | - | - | +++ | + | (+++) | - | (+++)<sup>2*</sup> |
|  | Enrichment | 0 | + | (+++) | + | (++) | + | +++ | + | (+++) | + | (++)<sup>2*</sup> |
|  |  | 7 | + | (+++) | + | (++) | + | +++ | + | (+++) | + | +++ |
| Arugula | Direct extraction | 0 | - | - | - | - | - | (++)<sup>3*</sup> | + | (++) | - | (+++)<sup>2*</sup> |
|  |  | 7 | - | - | - | - | - | (++)<sup>1*</sup> | + | (++) | - | (+++)<sup>1*</sup> |
|  | Enrichment | 0 | + | (+++) | - | - | + | +++ | + | (++) | + | +++ |
|  |  | 7 | + | (+++) | - | - | + | +++ | + | (++) | + | +++ |
| Cilantro | Direct extraction | 0 | - | - | - | - | - | - | (+)<sup>1*</sup> | (++) | (+)<sup>1*</sup> | (+++)<sup>3*</sup> |
|  |  | 7 | (+)<sup>1*</sup> | (++)<sup>2*</sup> | - | - | - | - | (+)<sup>3*</sup> | (+++) | (+)<sup>2*</sup> | (+++)<sup>3*</sup> |
|  | Enrichment | 0 | (+)<sup>2*</sup> | (+++)<sup>2*</sup> | (+)<sup>1*</sup> | (++)<sup>2*</sup> | - | - | (+)<sup>3*</sup> | (+++) | (+)<sup>2*</sup> | (+++)<sup>3*</sup> |
|  |  | 7 | (+)<sup>2*</sup> | (+++)<sup>2*</sup> | (+)<sup>2*</sup> | (++)<sup>2*</sup> | - | - | (+)<sup>3*</sup> | (+++)<sup>3*</sup> | (+)<sup>3*</sup> | (+++)<sup>3*</sup> |

(\*): Number of positive replicates, -: Not detected or no signal, +: positive (qPCR), (++) : Medium signal, (+++): Strong signal

## DISCUSSION

The present study showed that bacteria associated with produce can carry various plasmids that might serve as link between the environmental and the human gut microbiome. Although initially low in abundance, tet resistant *E. coli* were isolated from all produce samples purchased after none selective enrichment. Contamination of produce with *E. coli* strains can occur in the field by contaminated soil (organic fertilizers), exposure to contaminated irrigation water or post harvest (12, 13). In this study, tet resistant *E. coli* isolates were mostly isolated from cilantro that was purchased in four Asian supermarkets in two cities, followed by mixed ready to eat salad and arugula. This might be a hint that produce imported from Asia might be a hotspot for contamination with *E. coli* carrying multidrug resistance plasmids. A high proportion of the tet resistant *E. coli* isolates was also resistant to penicillins (AM and AMX) followed by trimethoprim. Although it is difficult to compare among studies because of different methodologies used for isolation and resistance testing, our results are in line with high resistance levels to penicillins and trimethoprim previously reported for *E. coli* from irrigation water and vegetables (18), ready to eat salads (1), and lettuce (2). In the present study, tet resistance was commonly conferred by *tet*(A), partly confirming previous studies reporting *tet*(A) and *tet*(B) genes as the most common tet resistance genes in *E. coli* and *Salmonella spp*. isolated from ready to eat vegetables (1, 32). The rapid dissemination of tetracycline resistance among bacteria has been related to the occurrence of *tet* resistance genes on transposons and conjugative plasmids (22, 23, 33), but also to the selective pressure, e.g. the use of antibiotics in animal husbandries and the spread of tet resistance genes via organic fertilizers (31).

Plasmid-mediated multidrug resistance plays an important role in the transfer of ARGs around the world (34). Our study showed that *E. coli* isolates from produce harbored various plasmids belonging to replicon types IncF, IncI1, IncX1, IncU, IncN, and IncHI1, with IncF plasmids being the most frequently detected. IncF plasmids were found predominantly in *E. coli* isolated from drinking water (35) and poultry farms (36). In our study IncFII was the most frequently detected (36.5%) replicon type, followed by IncFIB (15.9%), which is in line with studies on *E. coli* recovered from pigs and human (37), wastewater (38), and animals (39). The combination of replicon types IncFII and IncFIB in two isolates is consistent with a report on *Enterobacter cloacae* from lettuce (3). However, we cannot exclude that these replicons are located on the same plasmid, as several studies have reported the combination of replicon types as a multi-replicon on a single plasmid (40–42), likely due to co-integration (28). In this study, tet resistant *E. coli* isolates which carried IncF plasmids were positive also for *tet*(A), *aadA, sul1, sul2, sul3, qacE/qacEΔ1, qnrB, qnrS* or *bla*_TEM_ genes. Previous reports found that IncF plasmids can carry genes conferring resistance to all major antibiotic classes including aminoglycosides, β-lactams, phenicols, tetracyclines, sulfonamides, and fluoroquinolones (37, 39, 43).

The NHR IncI1 plasmid types were the second most dominant replicon type (34.9%) and Inc11 positive isolates also carried multiple ARGs. In this study, strains carrying Inc11 plasmids were also positive for class 1 integron integrase gene *intI1* and a diverse set of resistance genes, namely *tet*(A), *sul2, strA*, *bla*_TEM_, *qacE/qacEΔ1, aadA, sul3, qnrS*, and *merRTΔP*. In a recent study, IncI1 plasmids from irrigation water and lettuce carried genes *sul1*, *tet*(A), *aadA, strA*, and *bla*_TEM_ as well as *intI1* (18). Similar phenotype and genotype profiles among *E. coli* strains from the current study and those recovered in previous studies from clinical samples, the environment or other foods indicate that produce may play a potential role in the dispersal of *E. coli* carrying plasmid-localized ARGs. Thus, plasmids belonging to the IncF and IncI groups have the potential to be a major contributor worldwide to the propagation of ARGs within enteric bacteria. One dissemination route of enteric bacteria carrying IncF and IncI plasmids might be the consumption of produce.

The newly developed TaqMan probe-based RT-PCR assays demonstrated high specificity in detecting these plasmids *in E. coli* isolates and PCR positives were also assigned by PBRT which in addition enables sub-typing.

This is the first study identifying NHR plasmids such as IncX1 and IncHI1 and broad-host-range (BHR) plasmid IncU in *E. coli* isolates recovered from cilantro leaves. Interestingly, IncX plasmids were detected in *E. cloacae* from lettuce (3). IncHI1 plasmids were previously reported in *E. coli* and *Citrobacter youngae* isolates from water and healthy calves, respectively (44), while the first IncU plasmids were isolated from *Aeromonas_salmonicida* (45), and later from *Aeromonas caviae* from hospital effluent in the UK (46). In general, a low prevalence of ESBL-producing *E. coli* was found on produce, which is similar to previous studies (19, 47, 48). In the present study, the occurrence of ESBL-producing *E. coli* were only isolated from cilantro (2.8%).

To our knowledge, this is also the first report of *E. coli* isolates from cilantro that were positive for conjugative IncN plasmids, *bla*_CTX-M-1_, and resistance to 3^rd^ generation cephalosporins. The *bla*_CTX-M-1_ gene was also reported on plasmids belonging to the IncN family in *E. coli* from farm workers, animals, humans, and the environment (49–51). Although IncN plasmids are able to replicate in a variety of *Enterobacteriaceae*, they are most frequently found in *E. coli* and *Klebsiella pneumoniae*, where they contribute to the dissemination of cephalosporin and carbapenem resistance (52).

The results of the present study is investigation showed that *E. coli* isolates harboring the *bla*_CTX-M-1_ gene also conferred resistance to at least seven classes of antibiotics tested. Moreover, *E. coli* harboring CTX-M genes were recently reported from lettuce and irrigation water (4, 53), raw vegetables (32, 53), and coastal waters (54, 55). Kim et al. reported that ESBL-producing *E. coli* and *Klebsiella pneumoniae* carrying CTX-M were detected in ready to eat vegetables (56). A recent study has detected *bla*_TEM_ genes in association with IncF and IncI1 plasmids from irrigation water and lettuce (18).

In previous reports, the occurrence of *sul1* and *qacE/qacEΔ1* was frequently associated with class 1 integrons (3, 57). Unexpectedly, only 27% and 36.5% of *sul1*- and *qacE/qacEΔ1*- positive isolates carried the *intI1* gene, respectively, indicating that atypical class 1 integrons were more prevalent among the isolates as previously also reported by Amos et al. (58).

In the present study, transferable tet resistance plasmids were also directly captured from the produce microbiomes on day 0 but not on day 7 after purchase, and the highest transfer frequency was observed in mixed salad, followed by cilantro and arugula. Differences in observed frequencies of transconjugants could be due to different abundance of bacteria with conjugative plasmids in the various sample types, or due to real differences in the frequencies of plasmid transfer. The latter might be affected by the metabolic activity of the produce microbiome, as plasmid transfer frequency is known to depend not only on plasmid-specific characteristics, but also ecological factors affecting the metabolic activity of bacteria (59). Replicon types IncFII, IncFIB, IncI1, and IncP-1β were captured from cilantro leaves whereas only IncFII plasmids were captured from mixed salad. IncF (FII and FIB) plasmids were prevalent among tet resistant transconjugants from both produce. Most of the IncF plasmids exogenously captured harbored *bla*_TEM_, *tet*(A), and *qnrS* genes. One IncI1 plasmid was captured from cilantro and another one in combination with replicon type IncFIB. The conjugative plasmids carried *tet*(A) and *bla*_TEM_ genes. Finally, four IncP-1β plasmids were captured from cilantro leaves and two of them in combination with replicon type IncFII. IncP-1 plasmids have been frequently captured by exogenous plasmid isolation from various environments such as sewage sludge (60), manure (23), and water (61). However, the first isolations of IncP-1 plasmids were from clinical isolates (62, 63). The IncP-1β plasmids carried genes conferring resistances to antibiotics *tet*(A), *strA*, and *bla*_TEM_ but also mercury compounds (*merRTΔP*) and disinfectants (*qacE/qacEΔ1*).

In conclusion, this study showed that produce that we eat might contain bacteria such as *E. coli* carrying transferable multidrug resistance plasmids. Although *E. coli* numbers are typically low, our nonselective enrichments showed that proliferation can easily occur. Our study reports a specific TaqMan probe-based RT-PCR assays that can be used for rapid detection of IncF and IncI plasmids in *E. coli* isolates and exogenously captured plasmids as well as in TC-DNA extracted from enrichment cultures of leaves. However, quantifying these plasmids in TC-DNA directly extracted from the microbial fraction detached from leaves was impossible due to their low abundance in the microbiome, but IncF and IncI plasmids were detected in DNA extracted after previous enrichment. While these assays represent an important and useful tool to be implemented for monitoring the prevalence of IncF and IncI plasmids in isolates and the environment, negative results of these and other cultivation-independent methods can lead to an underestimation of the mobile resistome present in the rare microbiome of produce and other samples. This is the first study demonstrating that multi-drug resistance plasmids present in produce associated bacteria were transferable to sensitive *E. coli* recipients, a process that could occur in the human gut. The NHR plasmids IncF and IncI1 but also the BHR IncP-1β plasmids were captured from the produce. In particular the captured IncF and Inc1 plasmids conferred resistance towards several classes of antibiotics. Thus produce associated bacteria should be considered as important route of disseminating transferable antibiotic resistances which might be particularly relevant for patients under antibiotic treatment.

## MATERIALS AND METHODS

### Sample collection

A total of 24 samples from different produce (mixed salad, arugula, and cilantro) imported or locally produced was analyzed. The produce samples were purchased from different local supermarkets in Germany and collected in September-2016, June-2016 and Mai-2017, respectively. They were stored at fridge temperature and sampled on days 0 and 7, four replicates for each time point and produce type.

### Isolation and identification of tet resistant *E. coli*

For sampling, the produce was cut into smaller pieces using a sterile scalpel and mixed. Of each sample 25 g were placed in two stomacher bags (one for direct plating and the other for enrichment) and mixed three times with 75 ml buffered peptone water (BPW, Roth; Karlsruhe, Germany), with subsequent stomacher treatment performed with the Stomacher^®^ 400 (Seward Worthing, England) at high speed for 1 min. The enrichment cultures of fresh leaves in BPW were incubated at 37°C with shaking (150 rpm) for 18-24 h. In order to isolate tet resistant *E. coli*, appropriate dilutions of the sample suspensions and also of the enrichment cultures were plated on different culture media [Eosin Methylene Blue (EMB, Sifin; Berlin, Germany) and Chromocult coliform agar (CCA, Merck, Darmstadt, Germany)] supplemented with tetracycline (10 mg L^−1^). All plates were incubated at 37°C for 18-24 h. The presumptive *E. coli* colonies were picked from each sample and streaked onto EMB, CCA, and TBX chromogener Agar (TBX, Roth; Karlsruhe, Germany) for confirmation by colony morphology and further characterization. *E. coli* isolates were then confirmed by biochemical tests for indole production, methyl red and catalase activity (64). Furthermore, isolates were analyzed for the presence of the *gadA* gene (Glutamate Decarboxylase Genes) specific for *E. coli* using PCR (65). *E. coli* isolates were stored in Luria broth (LB, Roth; Karlsruhe, Germany) containing 15% glycerol at −80^°^C.

### Exogenous plasmid isolation

In order to capture tetracycline-resistance plasmids exogenous plasmid isolation via biparental mating was performed using [*Escherichia coli* CV601 gfp^+^, kanamycin (Km) and rifampicin (Rif) resistant; Heuer et al. (66)] as a recipient. The recipient strain was grown overnight in Tryptic Soy Broth (TSB; Merck, Darmstadt, Germany) supplemented with rifampicin (Rif) (50 mg L^−1^) and kanamycin (Km) (50 mg L^−1^). Two ml of the recipient strain culture was transferred into a sterile Eppendorf tube and centrifuged at 3,100 × *g* for 5 min and washed twice with 1:10 TSB. Then, the pellet was resuspended in 2 ml of 1:10 TSB. The bacterial suspensions (donor) of each sample on days 0 and 7 were prepared from enrichment cultures of fresh leaves as described above. 0.5 ml of recipient strain and 20 ml of each enrichment culture (donor) were mixed in a 50 ml falcon tube. As a background control, 5 ml of the enrichment cultures and 200 μl of the recipient were processed the same way as the samples. All mixtures were centrifuged at 3,100 × *g* for 10 min. The pellets were resuspended in 200 μl of 1:10 TSB and then spotted onto a filter for mating (Millipore filters, 0.22μm). Filters were incubated overnight at 28°C on Plate count agar plates (PCA; Merck, Darmstadt, Germany) supplemented with cycloheximide (Cyc) (100 mg L^−1^). After incubation, the filters were placed in 2 ml of sterile 0.85% NaCl solution in a 50 ml falcon tube. Each filter was washed by vortexing for 1 minute. Serial 10-fold dilutions were done and appropriate dilutions were plated on PCA agar supplemented with rifampicin (Rif 50 mg L^−1^), kanamycin (Km 50 mg L^−1^), cycloheximide (Cyc 100 mg L^−1^), and tetracycline (Tet 15 mg L^−1^) to select for tetracycline-resistant transconjugants. Background controls of bulk soil and the recipient controls were plated on the same selective media. Numbers of recipient cells were determined by applying three replicate 20 μl drops per each serial dilution (10^−5^ to 10^−8^) of all mating mixes on PCA with km (50 mg L^−1^), rif (50 mgL^−1^), and cyc (100 mg L^−1^). All plates were incubated at 28°C for up to 3 days. Transconjugants were determined by green fluorescence resulting from the green fluorescence protein (GFP). The identity of putative transconjugants was confirmed by BOX-PCR (67). Transfer frequencies were calculated as total number of transconjugants divided by the total number of recipients.

### Antibiotic susceptibility testing

Antimicrobial susceptibility testing was performed by the disk diffusion method on Müller-Hinton agar (MH; SIGMA-ALDRICH CHEMIE GmbH, USA), according to the EUCAST (European Committee on Antimicrobial Susceptibility Testing). The antibiotics (μg) (BD, Bacton, Dickinson and Company, USA) used in this study were: amoxicillin (25), ampicillin (10), cefotaxime (30), ceftazidime (30), ceftriaxone (30), chloramphenicol (30), ciprofloxacin (5), colistin (10), tet (30), doxycycline (30), streptomycin (10), gentamicin (10), ofloxacin (5), kanamycin (30), nalidxic acid (30), trimethoprim (5), sulfadiazine (250). Tet resistant *E. coli* isolates were streaked onto LB agar supplemented with tet (10 mg L^−1^), while tet resistant *E. coli* CV601 transconjugants were streaked on plate count agar plates (PCA, Merck, Darmstadt, Germany) supplemented with tet (15 mg L^−1^), km (50 mg L^−1^) and rif (50 mg L^−1^). *E. coli* CV601 strain was used as a negative control. The bacterial suspension was prepared from a single colony in normal saline (0.85% NaCl) to a density of 0.5 McFarland turbidity standard. Cotton swabs were used for streaking the suspension onto MH agar plates. After air drying, antibiotic discs were placed on the plates. Then all plates were incubated at 37°C for 18-24 h. The inhibition zone was measured. The results were interpreted according to the guidelines of EUCAST. Clinical and Laboratory Standards Institute (CLSI) recommendations were used when antibiotic breakpoints in EUCAST guidelines were absent (i.e. doxycycline, streptomycin, tetracycline, and nalidixic acid). ESBL production was confirmed among tet resistant *E. coli* isolates and transconjugants by double-disc diffusion test (DDT) (47). The phenotypic confirmatory test was confirmed as ESBL producer according to the CLSI.

### TC-DNA extraction

The bacterial fraction detached from fresh leaves directly or after an enrichment culture of each sample as described above were pelleted by centrifugation at 3,100 × *g* for 15 min at 4°C. Total community-DNA was extracted from the pellet using the FastDNA^®^SPIN Kit for soil (MP Biomedicals, Heidelberg, Germany), according to the manufacturer’s instructions. The quality of extracted DNA was determined by agarose gel electrophoresis. The extracted DNA was stored at −20°C until further analysis.

### Genomic DNA extraction

Genomic DNA was extracted from overnight cultures of tet resistant *E. coli* isolates, transconjugants and the recipient strain with Qiagen genomic DNA extraction kit (Qiagen, Hilden, Germany) using a silica-based kit (silica bead DNA extraction kit; Thermo Scientific, St. Leon-Rot, Germany). The extracted genomic DNA was stored at - 20°C until further analysis.

### Primer-probe design (IncF, IncI1, and IncI2 plasmids)

As it is known that relaxase genes can be used for classification of the mobilization systems of plasmids (68), the *traI* gene region was chosen as a target region to design primers detecting IncF, IncI1 and IncI2 plasmid sequences. A total of 4,530 plasmid DNA sequences were downloaded from NCBI (NCBI, Batch Entrez) using the 4,602 plasmid accession numbers found in GenBank by Shintani et al. (69), among which 298 plasmids were identified as MOBF. The CDS of the MOBF plasmids were aligned using tBLASTn against the relaxase TraI of the F plasmid (AP001918), resulting in 110 protein sequences sharing > 50% identity and >70% coverage. The 110 protein sequences closely related to TraI were aligned using MAFT multiple sequence alignment software version 1.3.3. The alignment produced was back translated using the EMBOSS Backtranseq tool and used to generate a set of degenerated primers and probes using Primer3. All those steps were carried out in Geneious^®^ 8.1.9. At best, 83 of the 110 *traI* nucleic acid sequences could be targeted by one set of designed primers and probe (see Table 4). Those sequences belonged mostly to plasmids isolated from *Salmonella enterica* and *Escherichia coli* and few from *Klebsiella pneumoniae* and *Shigella* spp. The plasmids corresponded to a part of the subclade MOBF12 defined by Garcillán-Barcia et al. (70), which is comprised of the phylogenetically broad IncF complex. When tested against the 4,530 plasmids the primers-probe targeted 92 plasmids in the database, 89 plasmids belonged to the MOBF group (298 plasmids recovered belonged to this group) and 3 were annotated as “non-mob” and belonged to any MOB group. When looking at the “Inc” classification, 73 of the targeted plasmids belonged to the IncF (including FI and FII), 14 IncZ and 5 to the pCD1 type. Among the 4,530 plasmid sequences 243 carried a *rep* gene belonging to the IncF group, indicating that the primer/probe cannot detect all possible IncF plasmids.

**TABLE 4.**
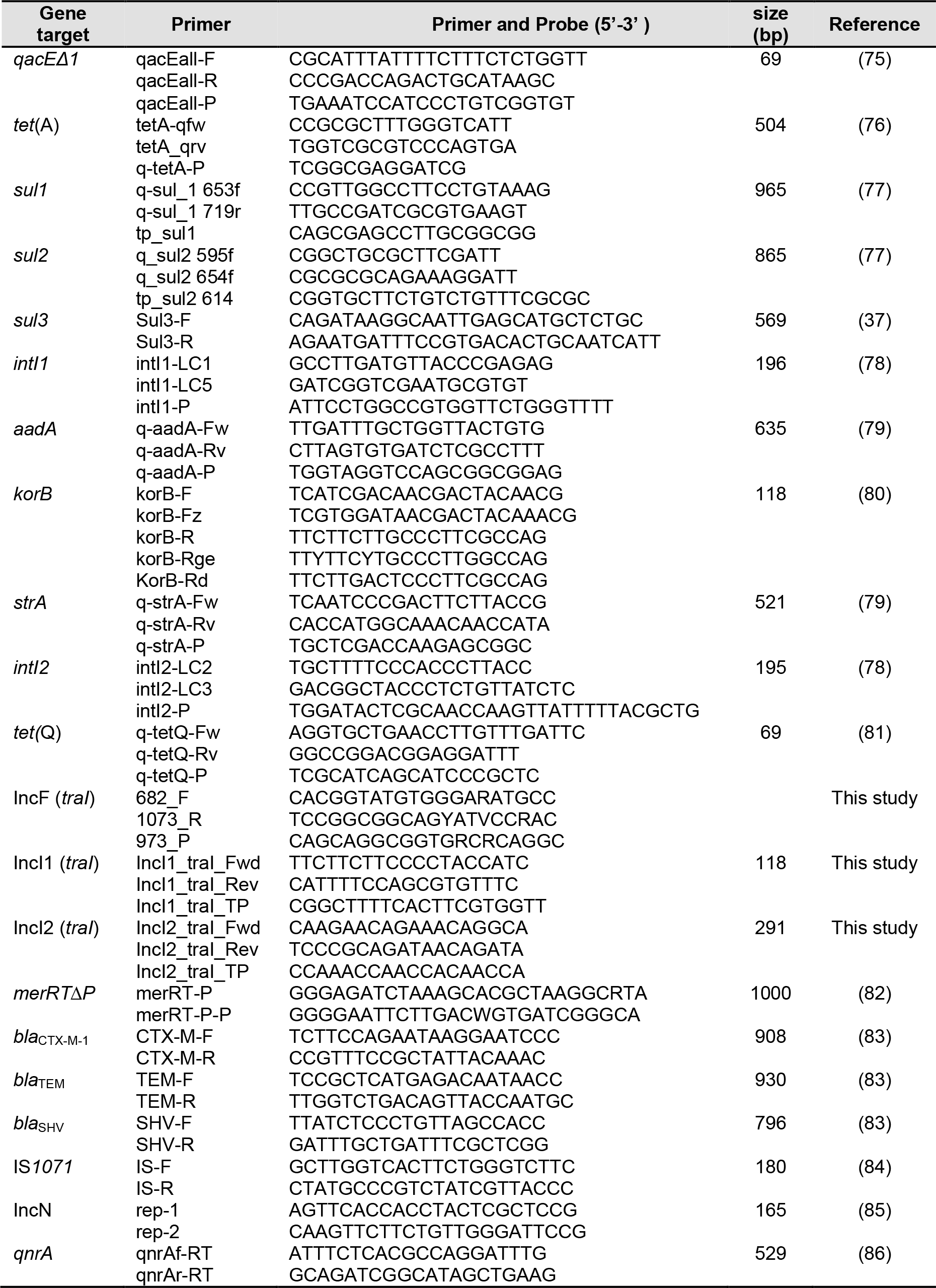
PCR and qPCR primer systems used in this study.

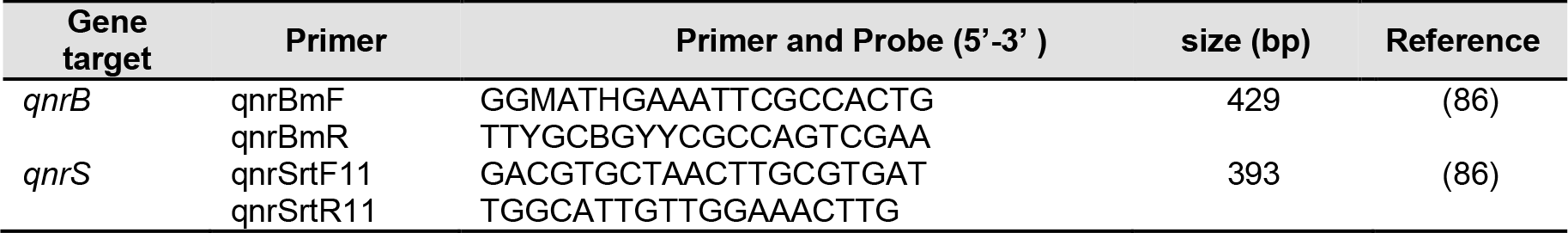

Available IncI plasmid sequences were downloaded from NCBI and the *traI* genes were aligned using the software CLC Main Workbench software version 8 (CLC bio, Qiagen^®^ company) with standard settings for alignments and primers were designed to match conserved regions of the *traI* gene (see Table 4). The specificity was confirmed in silico with NCBI primer BLAST and with a set of plasmids from other incompatibility groups. Plasmids used for this test were R388, pB10, pHHV216, RSF1010, pSM1890, RP4, pHH3-414, pHH2-227, pRA3, RN3, RSF1010, pTH10, pTP6, R751, pQKH54, pKS208, pEST4002, pJKJ5, pMCBF1, pRMS149, pCAR1, pD2RT, pD67, pWW0 and pST527 from which none was amplified.

### Detection of IncF, IncI plasmids by real-time PCR

The RT-PCR assay was performed under standard conditions, all qPCR reactions were set up in a 25 μL reaction volume using a Hot Start Taq DNA Polymerase (M0495L, New England BioLabs, Inc, UK) containing 5 μL of template DNA, 300 nM of primer (reverse), 50 nM of primer (forward), and 50 nM of probe for IncF, while 300 nM of each primer (forward and reverse) and TaqMan probe for IncI. All Primers and probes are described in Table 4. The following PCR program was used for amplification: 10 min at 95°C, followed by 40 cycles of 95°C for 30 sec and 60°C for 1 min. The assays were carried out in triplicate with real-time PCR 5’-nuclease assays (TaqMan qPCRRT-PCR) in a CFX96 real-time PCR detection system (Bio-Rad, Hercules, CA). Negative controls were included in all tests, and they consisted of all the elements of the reaction except for the template DNA.

Standard plasmids were used to construct a full standard curve in duplicate in each qPCR run. Standard plasmids were constructed by cloning the purified PCR products amplified from the plasmids R64 for IncI1, pHNSHP45 for IncI2, and IncF plasmids for the sub_MOBF12 using the corresponding primer pairs used for the RT-PCR, into TransforMax™ EC100™ Electrocompetent *E. coli* (Epicentre), in pJET1.2 using the Thermo Scientific CloneJET PCR Cloning Kit (Thermo Scientific).

### Detection of target genes by real-time qPCR and PCR

The target genes in genomic DNA extracted from tet resistant *E. coli* isolates and transconjugants were detected by RT-PCR 5’-nuclease assays (TaqMan or EvaGreen RT-PCR) in a CFX96 RT-PCR detection system (Bio-Rad, Hercules, CA) or by PCR for class 1 and 2 integrons integrase genes [*intI1* and *intI2*], *korB* (IncP-1 plasmids), *qacE/qacEΔ1* encoding quaternary ammonium compound resistance, [*aadA* and *strA*] encoding streptomycin and spectinomycin resistance, tetracycline resistance genes [*tet*(Q) and, *tet*(A)], *merRTΔP* gene part of the mercury resistance operon, [*sul1*, *sul2* and *sul3*] encoding sulfonamide resistance, [*qnrA, qnrB, qnrS*] encoding fluoroguinolone resistance, β-lactam resistance genes [*bla*_TEM_, *bla*_SHV_ and *bla*_CTX-M-1_], IncN (*rep*), and Insertion Sequence IS *1071* (represented by *tnpA* gene). The DNA of the recipient strain *E. coli* CV601 was included as negative control. The primers and probes targeting these genes and PCR and qPCR conditions are listed in (Table 4).

### Conjugation assay

Tet resistant *E. coli* isolates positive for ESBL (EK2.29, EK3.43, and EK3.44) were examined for their ability to transfer resistance. Briefly, the donors and the rifampicin and kanamycin resistant *E. coli* CV601 recipient strain were grown in LB broth overnight at 37°C. 500 μl of overnight cultures of each donor and recipient strains were mixed in 1 ml LB broth, and incubated at 37°C for 24 h without shaking. 100 μl of the conjugal mixture was spread on LB agar (Roth; Karlsruhe, Germany) containing Rif (50 mg L^−1^), Kan (50 mg L^−1^), CTX (2 mg L^−1^), and Tet (15 mg L^−1^), and incubated at 37°C for 24-48 h. The transconjugants were verified by BOX-PCR and further tested for the antibiotic resistance phenotypes and genotypes as described above.

### Plasmid DNA extraction and detection by Southern blot hybridization

Plasmid DNA extraction from tet resistant transconjugants (pBC1.1, pBC1.3, pBC1.9, and pBC1.12) captured exogenously from the enrichment of cilantro leaves was performed using Qiagen Plasmid Mini Kit (Qiagen Inc., Hilden, Germany) according to the manufacturer’s instructions. In order to detect the IncP-1 plasmids with Southern blot hybridization in these transconjugants, plasmid DNA was digested with the restriction enzyme *NotI* (Thermo Fisher Scientific, Waltham, MA, USA), and fragments were separated by electrophoresis on a 1% agarose-TBE gel as described previously (71). Southern blot hybridization was preformed with digoxigenin (DIG) labeled probes generated from PCR amplicons which were obtained from reference plasmids R751 for IncP1-β as previously described by Binh et al. (72).

### Plasmid replicon typing

PBRT was used to identify the incompatibility group of plasmids in tet resistant *E. coli* and transconjugants, and to confirm the presence of IncF and IncI plasmids as determined via the newly developed RT-PCR method as described above. This was done by PCR amplification on genomic DNA of the strains using primer sets for 30 replicons: HI1, HI2, I1, I2, X1, X2, X3, X4, L, M, N, FIA, FIB, FIC, FII, FIIS, FIIK, FIB KN, FIB KQ, W, Y, P1, A/C, T, K, U, R, B/O, HIB-M, and FIB-M, representative of major plasmid incompatibility groups among Enterobacteriaceae (73, 28). PCR products were separated by electrophoresis on a 2.5% agarose-TBE gel and stained with ethidium bromide.

### Detection of IncF and IncI plasmids, *tet*(A) and class 1 integrase gene *intl1* via PCR-Southern blot hybridization and RT-PCR in TC-DNA

PCR-Southern blot hybridization or RT-PCR was used to detect *tet*(A), *intI1*, IncF, and IncI plasmids in TC-DNA extracted from microbial fraction detached from leaves directly or after an enrichment step on days 0 and 7. The PCR products were separated on a 1% agarose-TBE gel electrophoresis and then transferred to a positively charged nylon membrane (GE Healthcare, UK). Southern blot hybridization was carried out with digoxigenin (DIG) labeled probes generated from PCR amplicons which were obtained from reference plasmids pKJK5 for *intI1* and *tet*(A) as described by Dealtry et al. (74), and R64 for IncI1, pHNSHP45 for IncI2, and IncF plasmids. The primers and PCR conditions are listed in Table 4.

## ACKNOWLEDGMENTS

The authors are thankful to Ute Zimmerling and Lena Rauch for their technical assistance. K.B. had a scholarship from the Libyan government, S.J. was funded by the German Environment Agency (Umweltbundesamt) (FKZ 3713 63402), and T.S. was funded by the United States Department of Agriculture (USDA) NIFA award # 2018-67017-27630.

